# Detection of *Xanthomonas translucens* pv. *undulosa* from wheat by quantitative PCR

**DOI:** 10.1101/2021.06.10.447895

**Authors:** Alian Sarkes, Yalong Yang, Snezana Dijanovic, Michael W. Harding, David Feindel, Jie Feng

## Abstract

A probe-based quantitative PCR (qPCR) protocol was developed for detection and evaluation of the wheat bacterial leaf streak pathogen *Xanthomonas translucens* pv. *undulosa* (Xtu). The protocol can also detect *X. translucens* pv. *translucens* (Xtt), but can’t differentiate the two pathovars. When tested on DNA from plant, non-target bacteria and culture of microorganisms from wheat seeds, the qPCR showed a high specificity. On purified Xtu DNA, the qPCR was more sensitive than a loop-mediated isothermal amplification (LAMP) assay. When DNA samples from a set of serial dilutions of Xtu cells were tested, the qPCR method could repeatedly generate quantification cycle (Cq) values from the dilutions containing ≥1,000 cells. Since 2 µL of the total of 50 µL DNA was used in one reaction, one qPCR reaction could detect the presence of the bacteria in samples containing as few as 40 bacterial cells. The qPCR could detect the bacteria from both infected seed and leaf tissues. For seed testing, a protocol for template preparation was standardized, which allowed one qPCR reaction to test DNA from the surface of one seed. Thus, the qPCR system could theoretically detect Xtu and/or Xtt in samples where the bacteria had an average concentration ≥40 cells per seed.

## Introduction

*Xanthomonas translucens* is a species of gram-negative bacteria causing bacterial leaf streak (BLS), also known as bacterial leaf stripe and black chaff, on cereals (Sapkota et al. 2020). The disease was first reported on barley (Jones et al. 1917) and wheat (Smith et al. 1919) in the USA and the pathogens were named *Bacterium translucens* and *B. translucens* var. *undulosum*, respectively. Once Dowson (1939) created the genus *Xanthomonas, Bacterium translucens* was reclassified as *X. translucens*. BLS has been found in almost all wheat- and barley-growing areas and can cause significant yield loss and reduction in seed quality (Sapkota et al. 2020). In Canada, BLS on wheat and barley was found in Manitoba as early as in 1922 (Drayton 1923).

The taxonomy and nomenclature of *Xanthomonas* has been very confusing and has undergone many changes driven by new studies on the bacterial biology, phylogenetics and genomics (Sapkota et al. 2020). Strains of *X. translucens* causing diseases on small grains were classified into a group called translucens group (Bragard et al. 1995; Vauterin et al. 1995). The translucens group was believed to contain three true biological entities, pathovars (pv) *cerealis, translucens* and *undulosa*, with pv. *translucens* pathogenic to barley, pv. *undulosa* pathogenic to both barley and wheat, and pv. *cerealis* pathogenic to barley, wheat, oat and bromegrass (Sapkota et al. 2020).

An effective disease management program is dependent on timely and proper identification to the causal agent of the disease. Due to the complicated subspecies structure of *X. translucens*, diagnosis into the pv level has been difficult. PCR and quantitative PCR (qPCR) protocols are not available. There is one loop-mediated isothermal amplification (LAMP) protocol (Langlois et al. 2017), which can specifically detect pv. *undulosa, translucens* and *secalis* but can’t differentiate the three pathovars.

BLS has been increasing in frequency and severity in the wheat-producing areas of the USA (Adhikari et al. 2020). This is also true in Western Canada, and recently the causal agent was confirmed to be Xtu (Tambong et al. 2021). Since 2019, the Alberta Plant Health Lab (APHL) has received grain samples from wheat plants with BLS symptoms. The frequency of samples increased in 2020. The LAMP protocol was very useful for detection and confirmation of Xtu and/or Xtt, but it could not discriminate between them, nor provide quantitative results. The purpose of this study was to develop a qPCR-based diagnostic protocol for detection and quantification of the BLS pathogens, and attempt to improve sensitivity, specificity and quantification in grain and leaf tissues. Herein we report the development of the qPCR system and the results obtained from evaluating the efficiency of this system.

## Materials and Methods

### Chemicals and standard techniques

All chemicals and instrument were purchased from Fisher Scientific Canada (Ottawa, ON) unless otherwise specified. All primers and probes were synthesized by Integrated DNA Technologies (Coralville, IA). PCR and LAMP were performed in a ProFlex PCR system. qPCR was performed in a CFX96 touch real-time PCR detection system (Bio-Rad Canada, Mississauga, ON).

### *Xanthomonas translucens* pv. *undulosa* isolates

Three *X. translucens* pv. *undulosa* (Xtu) isolates (Fig. 1a), named Xtu1, Xtu2 and Xtu3 were isolated from three wheat seed samples. These three samples were submitted to the APHL from different wheat fields with observed BLS symptoms in one county during the 2020 growing season. Upon receiving, seeds were cut into halves and incubated on Wilbrink’s boric acid-cephalexin medium (WBC) plate (Duveiller 1994) at 28°C in the dark for two days. Any bacteria growing on the media around the seed were purified on 2.3% (w/v) nutrient agar by the streak plate method. The Xtu isolates were identified based on morphologies, by fulfilling Koch’s postulates with artificial inoculation, symptomization (Fig. 1b) and re-isolation, and by DNA barcoding of the 16S ribosomal RNA and the chaperonin *cpn60* gene (GenBank accession numbers: MW784564-MW784566 for 16S and MW759847-MW759849 for *cpn60*). PCR amplification of the 16S and *cpn60* followed Klindworth et al. (2013) with the primer pair S-17/A-21 and Sahin et al. (2010) with the primer pair H1594/H1595, respectively. PCR product was purified by the Promega Wizard SV Gel and PCR Clean-Up System and sequenced by University of Alberta (Edmonton, AB). The three isolates were maintained on 2.3% (w/v) nutrient agar in darkness at 28°C. From 48-h nutrient agar cultures, cell suspensions were prepared. Cell concentration in suspensions were measured by OD600 using a NanoDrop 1000 spectrophotometer and by direct count using a haemocytometer.

**Fig. 1.**
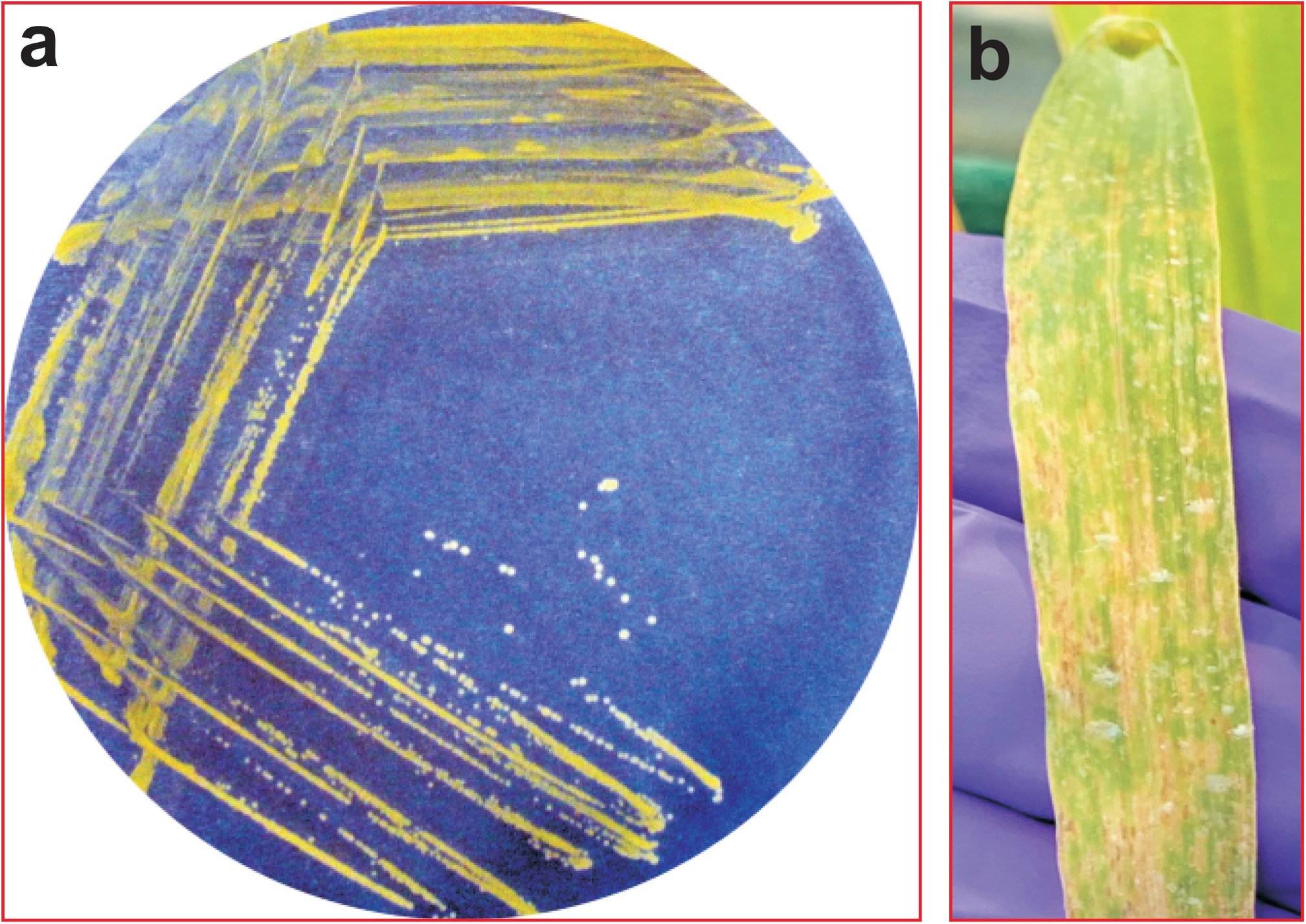
Morphologies of *X. translucens* pv. *undulosa* and the disease symptoms. (a) A *X. translucens* pv. *undulosa* isolate on nutrient agar plate, which was isolated from an infected wheat seed. (b) A symptomized wheat leaf after inoculated with this isolate.

### DNA extraction

All DNA extraction was conducted using the DNeasy Plant Pro kits (Qiagen Canada, Toronto, ON) manually unless otherwise specified when a Qiacube (Qiagen Canada) was used. DNA extracted from each sample was eluted in 50 µL water (manual) or elution buffer EB (Qiacube). For DNA extraction from bacteria, 1 or 1.5 mL cell suspension adjusted to the required concentration was centrifuged at 5,000 rcf for 5 min and then the supernatant was removed to leave 100 µL remaining to avoid disturbing the cell pellet. DNA was extracted from the resultant 100-µL samples.

### Primer design

When this study was initiated, there were three *X. translucens* strains with whole genome sequences (wgs) available in the National Center for Biotechnology Information (NCBI) database: ICMP11055 (pv. *undulosa*), XT4699 (pv. *undulosa*) and DSM 18974 (pv. *translucens*) (Sapkota et al. 2020). Among the wgs, only that of pv. *translucens* strain DSM 18974 was fully annotated (Accession number LT604072; checked June, 2021). In the annotations, in addition to genes encoding known proteins or proteins with homolog(s) in other species, there were 276 genes without any predicted function, encoding 165 “hypothetical proteins”, 39 “hypothetical secreted proteins” and 72 “unnamed protein products”. Of these 276 genes, those with a DNA sequence ≥200 base pairs (bp) were analyzed by Blastn against the NCBI nr/nt database. Twenty seven genes, consisting of genes for 16 hypothetical proteins, three hypothetical secreted proteins and eight unnamed protein products, produced Blastn hits exclusively in *X. translucens* pv. *translucens*, pv. *undulosa* and/or pv. *cerealis*. Based on the sequence of each of these 27 genes, a qPCR primer pair was designed using Primer-Blast (https://www.ncbi.nlm.nih.gov/tools/primer-blast/), with the following parameters differing from the default: PCR product size at 70-110 bp; primer melting temperatures (Tm) min. at 59°C, opt. at 62°C and max. at 65°C; data base: nr; organism: *Xanthomonas* group (taxid: 32033). The sequences of these primers are listed in Table S1. After the preliminary primer specificity test by SYBR Green-based qPCR, a probe named P11 was designed for the primer pair F11/R11 using Primer3 (https://bioinfo.ut.ee/primer3-0.4.0). The sequences of this primers/probe system are: F11, AATTGCACTGGCTGTCGCTG; R11, GGCAAGCGCCAAGAAGGAAA; P11, 56-FAM/GGCGTGGAC/ZEN/GTCTCTGTCCG/3IABkFQ.

### Primer sequence analysis

All 54 primers (Table S1) were further analyzed by Blastn against the NCBI nr/nt database. For each primer, the presence of identical sequence (only one alignment, with 100% coverage and 100% identity) in *X. translucens* pv. *translucens*, pv. *undulosa* and pv. *cerealis*, and other organism(s) was recorded (Table S1). Any primer pair with both the forward and reverse primers having identical sequences in organism(s) other than the three *X. translucens* pv. were removed from the primer list for further analysis. This ended up with 22 primer pairs (Table 1).

**Table 1.**
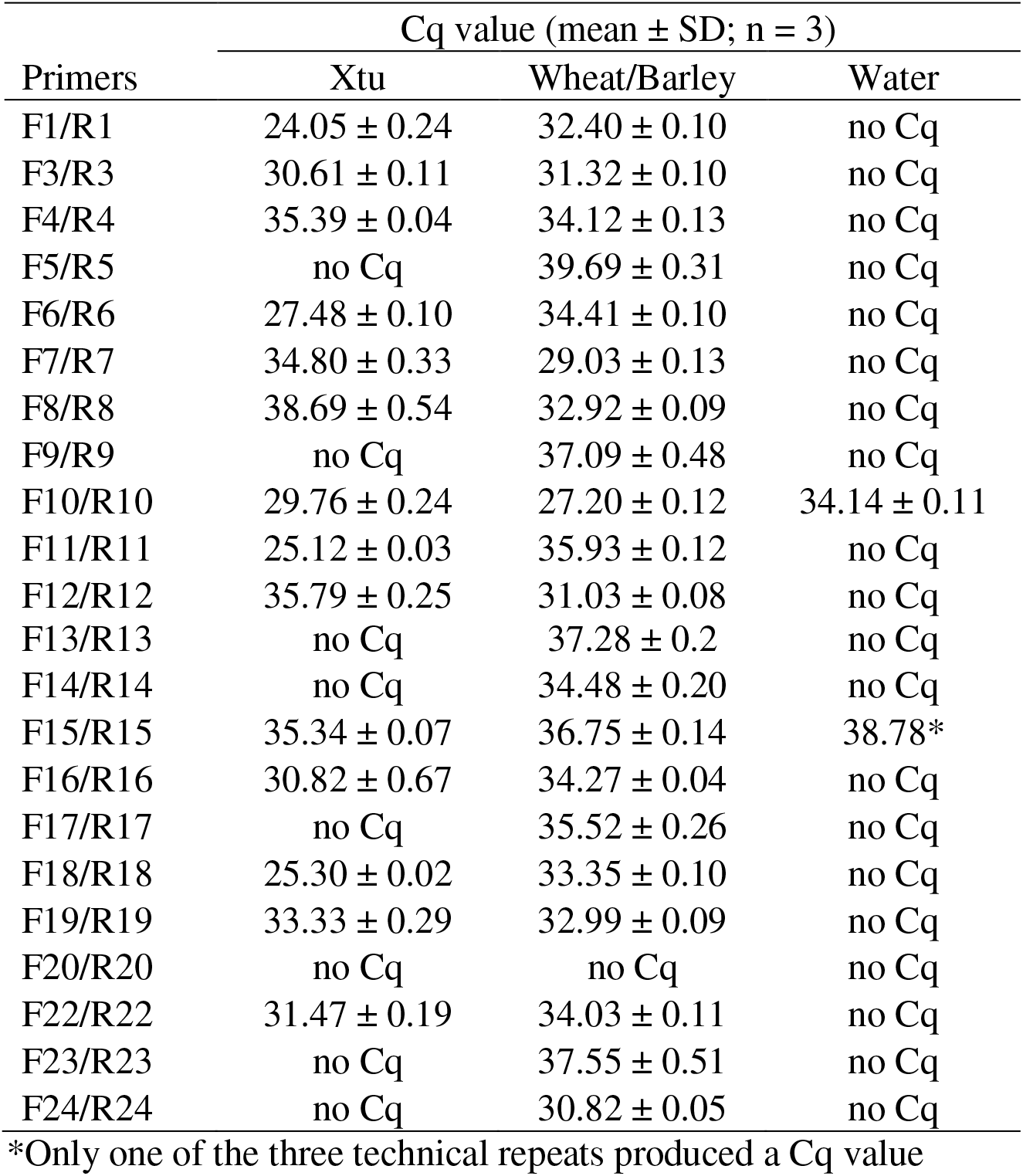
Screening on the specificity of 22 primer pairs by SYBR Green-based qCPR using DNA from *X. translucens* pv. *undulosa* (Xtu) and mixture of wheat and barley

### qPCR reaction

Each qPCR reaction was 25 µL containing 2 µL template and 0.25 µM of each primer. The probe-based qPCR reaction also contained 0.15 µM probe. SYBR Green-based qPCR and probe-based qPCR were conducted in SsoAdvanced universal SYBR Green supermix (Bio-Rad Canada) and PrimeTime gene expression master mix (Integrated DNA Technologies), respectively. The qPCR program consisted of an initial denaturation step of 95°C for 2 min, followed by 40 cycles of 5 s at 95°C and 30 s at 60°C. Each qPCR reaction was conducted with three technical repeats. In the specificity test of the 22 primer pairs by SYBR Green-based qPCR, a mixture of qPCR master mix, DNA template and water was prepared and aliquoted to reaction tubes, to which primers were added individually. In all other qPCR tests, mixture of qPCR master mix, primers/probe and water was prepared and aliquoted to reaction tubes, to which templates were added individually.

### LAMP reaction

The LAMP primer set Xt-CLS (Langlois et al. 2017) was used for confirmation of the qPCR test. All LAMP reactions were conducted in the WarmStart colorimetric LAMP master mix (NEB Canada, Whitby, ON). Each reaction was 25 µL in total and contained 2 µL template. Quantities of other components in each reaction followed the NEB’s instructions for the master mix. All reactions were conducted in individual 200-µL PCR tubes. The reaction program consisted of only one step in which the tubes were incubated at 65°C for 30 minutes. After the incubation, the reactions were checked visually and the results were recorded by photography.

### Primer screening by SYBR green-based qPCR

The 22 primer pairs were screened for specificity by SYBR green-based qPCR against two DNA samples: 1) mixed DNA (1:1:1) from the three Xtu isolates each at a concentration of 0.3 ng/µL and 2) mixed DNA (1:1) from leaves of 7-day old wheat (cv. CDC Teal) and barley (unknown cv.) seedlings generated on moist filter paper, each at a concentration of 30 ng/µL. After the screening, one primer pair (F11/R11) was selected for further analyses.

### Specificity test of the primer pair F11/R11 and primers with the probe F11/R11/P11

The specificity of the primer pair F11/R11 and primers with probe (F11/R11/P11) was tested by SYBR green-based qPCR and probe-based qPCR, respectively, on the following templates: mixed DNA (1:1:1) from the three Xtu isolates each at 3 ng/µL, DNA each at 30 ng/µL from wheat leaves and barley leaves of 7-day old seedlings, DNA at 3 ng/µL from pellets of 50 mL of YPG media (1% yeast extract, 1% peptone, 2% glucose, w/v) after incubating ten wheat seeds at 150 rpm 30°C for 48 hours (the pellets were prepared by centrifuging the culture after the seeds being removed at 4,000 rcf for 5 min) and DNA each at 3 ng/µL from five bacterial cultures on either 4% (w/v) PDA or 2.3% (w/v) nutrient agar including *Pseudomonas syringae* pv. *atrofaciens, P. syringae* pv. *syringae, Erwinia rhapontici, Clavibacter michiganensis* and *Xanthomonas campestris* pv. *pelargonii*. The five bacterial species were retrieved from the plant pathogen culture collection of APHL and their identities were confirmed by sequencing of the 16S ribosomal RNA.

### Sensitivity test of the primer pair F11/R11 and primers with the probe F11/R11/P11

Cell suspension was prepared from each of the three Xtu isolates. The suspensions were adjusted with water to OD600 = 0.6, which was equivalent to 1 × 10^9^ cells/mL, as confirmed by counting of the 100× and 1000× dilutions using a haemocytometer. One mL suspension of each of the three isolates was mixed in one tube. From 1.5 mL of the mixture, DNA was extracted and from which a set of 10× DNA serial dilution were prepared from 30 ng/µL to 30 fg/µL. In addition, from the 1 × 10^9^ cells/mL mixture a set of 10× cell serial dilutions were prepared until 1 × 10^3^ cells/mL. DNA was extracted from 1 mL of each dilution. All DNA samples were tested by LAMP, SYBR Green-based qPCR and probe-base qPCR. Furthermore, suspension of each of the three Xtu isolates was prepared from a culture plate alternative to the one used above. From each suspension, three 1-mL samples at a concentration of 100 cells/mL were prepared. DNA was extracted from these samples using a Qiacube to increase the consistency of DNA quantities from repeated cell samples. All DNA samples were tested by the probe-base qPCR.

### Test on infected wheat tissues

The probe-based qPCR and LAMP were conducted to test the three wheat grain samples from which the three Xtu isolates were isolated. From each sample, 100 seeds were soaked in 4 mL water in a 50-mL tube. The tube was vortexed for 30 sec and then from which three 1-mL samples of supernatant were prepared in 1.5-mL tubes. The supernatant samples were concentrated to 100 µL by centrifuging the tube at 11,600 rcf for 2 min and from which DNA was extracted, which is the DNA from the surface of 25 seeds. The 100 seeds were washed with 40 mL water three times and then soaked in water at a total volume of 10 mL in 4°C overnight. The 10-mL seed/water mixture was ground with a mortar and pestle and from the resultant slurry 100-µL samples were subjected to DNA extraction, which represents the DNA from one seed. In addition, DNA from infected wheat leaves was also tested by LAMP and qPCR. To prepare the infected leaves, 14-day old seedlings of wheat cv. CDC Teal were inoculated with Xtu1 in a inoculation chamber maintained at 90% relative humidity, 24°C/18°C (day/night) and a 16-h photoperiod. The inoculum was prepared at 1 × 10^8^ cells/mL in water supplemented with 2.5 g/L sand (Cat: S5631; Sigma-Aldrich Canada, Oakville, ON). Inoculation was conducted with a spray bottle 10 cm away from the plants until run-off. DNA was extracted from 100-mg samples of symptomized leaves at the 14^th^ day after inoculation (dai).

### Statistics

In all qPCR, means of the quantification cycle (Cq) values from the three repeats were calculated and treated as one data point. The qPCR standard curve was constructed by regression analysis using the SAS software (version 9.4; SAS Institute, Cary, NC). qPCR primer efficiencies were calculated as E = -1+10^(−1/slope)^ (Svec et al. 2015). All experiments with data shown in this paper were conducted at least twice with similar results.

## Results

### Primer screening

Twenty-seven qPCR primer pairs were designed each targeting a gene exclusively present in *X. translucens* pv. *translucens*, pv. *undulosa* and/or pv. *cerealis*. Blast the primer sequences against the NCBI database indicated that identical sequences of both the forward and the reverse primers in primer pairs F10/R10 and F11/R11 were only present in pv. *translucens* and pv. *undulosa* (Table S1). Therefore, these two primers pairs were potentially more specific than others.

Twenty-two primer pairs, including F10/R10 and F11/R11, were tested for specificity by SYBR Green-based qPCR (Table 1). Cq values were produced from wheat and barley DNA by most primers pairs. However, with the consideration that a specific primer pair should have low Cq from target and high Cq from non-target, F11/R11 was the best among the 22 primer pairs tested. This primer pair targets the GenBank accession number SCB06003, which is indicated as a hypothetical protein in the annotation of the wgs of the *translucens* DSM 18974 isolate peng1 (GenBank accession number: LT604072) and coded by an open reading frame at 210 bp. A Blast of the 210-bp sequence against NCBI wgs database (conducted June 2021) indicated that this gene is present in four pv. *translucens* strains with identical sequences (one from Germany, three from China) and ten pv. *undulosa* strains also with identical sequences (four from USA, six from China). The sequences in pv. *translucens* and pv. *undulosa* differed on three nucleotides (Fig. 2).

**Fig. 2.**
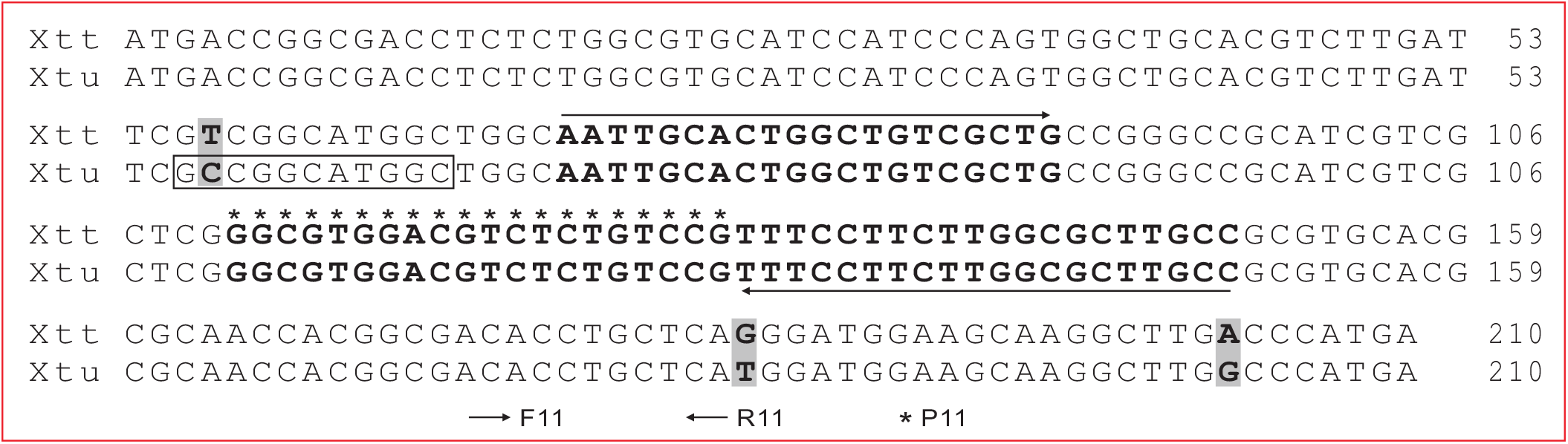
The sequences of the qPCR target in *X. translucens* pv. *translucens* (Xtt) and pv. *undulosa* (Xtu). Sequences of primers and probe are bold and indicated. The three single nucleotide differences between Xtt and Xtu are highlighted. The *Bgl* I restriction enzyme cutting site is framed.

### Specificity of F11/R11

Using DNA from five bacterial species, wheat, barley and YPG culture of wheat seeds, the specificity of F11/R11 was tested by SYBR Green-based qPCR and probe-based qPCR. From *P. syringae* pv. *syringae* and *P. syringae* pv. *atrofaciens*, one of the three technical repeats of the probe-based qPCR produced signal but with high Cq values (Table 2). No signal was produced from all other non-target DNA. In contrast, SYBR Green-based qPCR produced signals from all DNA templates, although the Cq of non-targets were higher than those of the target. These results indicated that the probe-based qPCR is more specific than the SYBR Green-based qPCR and the former is well-suited for Xtu diagnosis in suspicious wheat tissue samples.

**Table 2.** Specificity test of F11/R11/P11

| DNA template | Cq value (mean ± SD; n = 3) | |
| --- | --- | --- |
|  | Probe-based qPCR | SYBR Green-based qPCR |
| *Pseudomonas syringae* pv. *syringae* | 39.0* | 35.3 ± 0.2 |
| *P. syringae* pv. *atrofaciens* | 38.5* | 35.1 ± 0.2 |
| *Erwinia rhapontici* | no Cq | 34.9 ± 0.2 |
| *Clavibacter michiganensis* | no Cq | 38.1 ± 0.8 |
| *Xanthomonas campestris* pv. *pelargonii* | no Cq | 37.2 ± 0.9 |
| Wheat culture in YPG medium | no Cq | 31.3 ± 0.1 |
| Wheat | no Cq | 35.0 ± 0.1 |
| Barley | no Cq | 35.5 ± 0.1 |
| Mix of the three Xtu isolates | 21.5 ± 0.1 | 21.8 ± 0.2 |

### Sensitivity of F11/R11

The sensitivity of the probe-based qPCR was tested against two set of DNA: a set of serial dilutions of the Xtu DNA and DNA from a set of serial dilutions of Xtu cells. From the first set of DNA, the constructed standard curve had a slope of -3.350 and a R^2^ of 0.9996 (Fig. 3a). The efficiency of the qPCR was 0.99. A qPCR reaction could detect 10^1.78^ = 60 fg bacterial DNA (2 µL template taken from 30 fg/µL DNA sample), with a mean Cq value of 38.2. On the same set of DNA samples, LAMP could produce positive signal at ≥ 10^5.78^ fg (600 pg) DNA per reaction (Fig. 3b). These results indicated that the probe-based qPCR is 10^4^ times more sensitive than LAMP. From the second set of DNA samples, the constructed qPCR standard curve had a slope of -2.791 and a R^2^ of 0.9929 (Fig. 3c). The efficiency of the qPCR was 1.28. A qPCR reaction could detect DNA from 10^1.6^ = 40 bacterial cells (2 µL template taken from 50 µL DNA extracted from 1,000 cells). DNA was extracted by Qiacube from nine 100-cell samples, with three from each of the three Xtu isolates. When these nine DNA samples were tested by qPCR, most of the technical repeats failed to produce a Cq value (data not shown). The few that did produce a value had a Cq between 38.5 and 39.9. These results indicated that the qPCR system, which includes the DNA extraction, the qPCR program and reaction constitution, could detect the bacteria from a sample containing ≥ 1,000 bacterial cells.

**Fig. 3.**
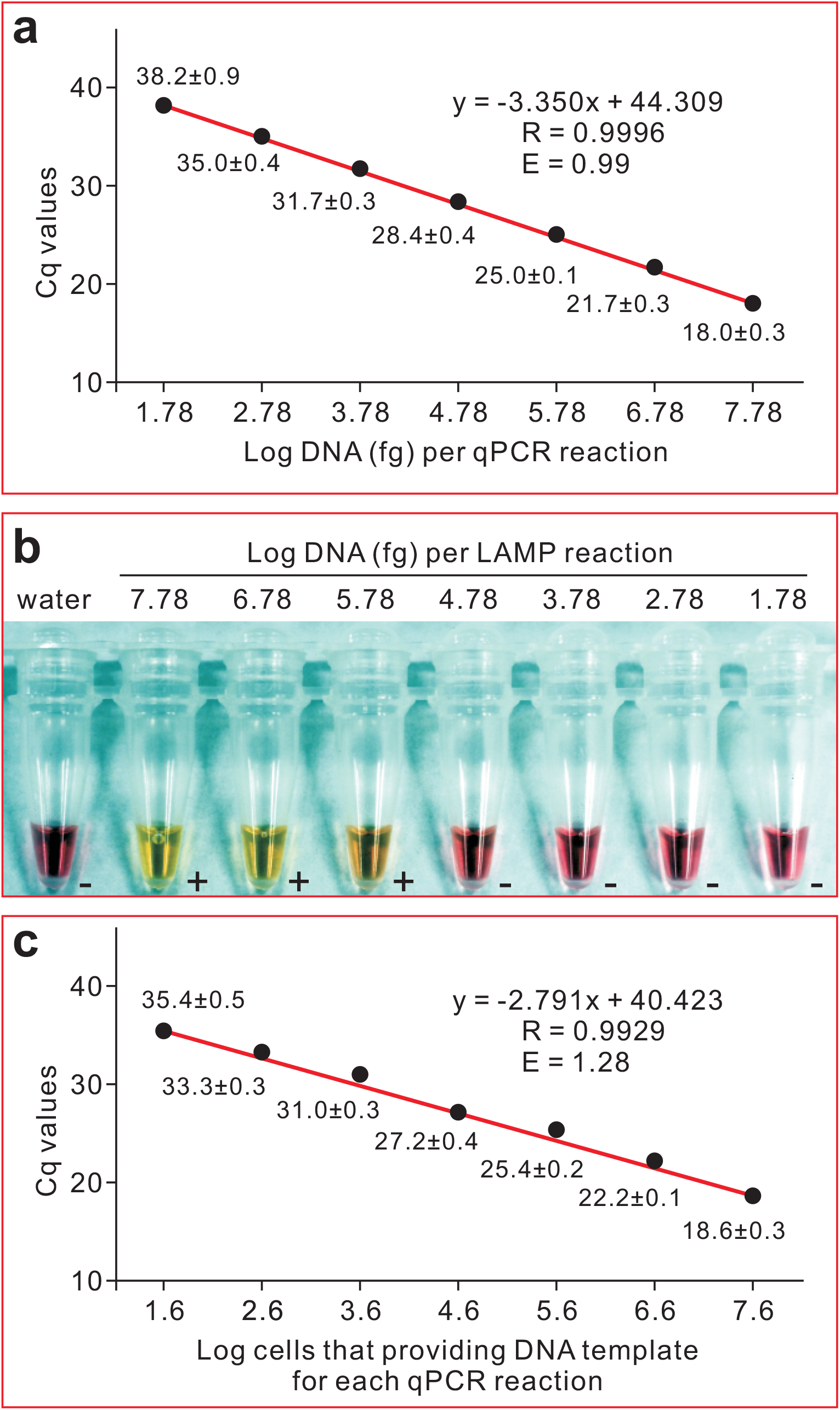
Sensitivity of the qPCR system and LAMP on the detection of *X. translucens* pv. *undulosa*. (a) and (c), qPCR standard curves generated from the mean of quantification cycle (Cq) values against log10 of template DNA in the reaction. (b) Sensitivity of the LAMP assay against log10 of template DNA in the reaction. The templates in (a) and (b) were a set of 10-fold serial dilutions of the Xtu DNA. The templates in (c) were DNA from a set of 10-fold serial dilutions of Xtu cells. In (a) and (c), the R^2^ score of the equation and the efficiency of the primers (E), are indicated over the curve. Efficiency was calculated as E = -1+10^(−1/slope)^. Each data point is shown as mean ± standard deviation (n=3).

### Detection of Xtu from plant samples

Three wheat seed samples were tested by LAMP and the probe-based qPCR. For all the three samples, LAMP could detect the presence of Xtu from surface of seeds but not from the washed seeds (Fig. 4a). In contrast, qPCR could detect the bacteria from both surface of seeds and washed seeds. These results indicated that the qPCR system and the sample preparation protocol are well-suited for seed testing. When serial dilutions of DNA samples from two infected wheat leaves were tested, qPCR again had higher sensitivity compared to LAMP. LAMP could not produce positive signal from DNA samples with a qPCR Cq value ≥ 29.4 (Fig. 4b). These results indicated that the qPCR system could also be used for detection/confirmation of Xtu within infected wheat leaf tissues.

**Fig 4.**
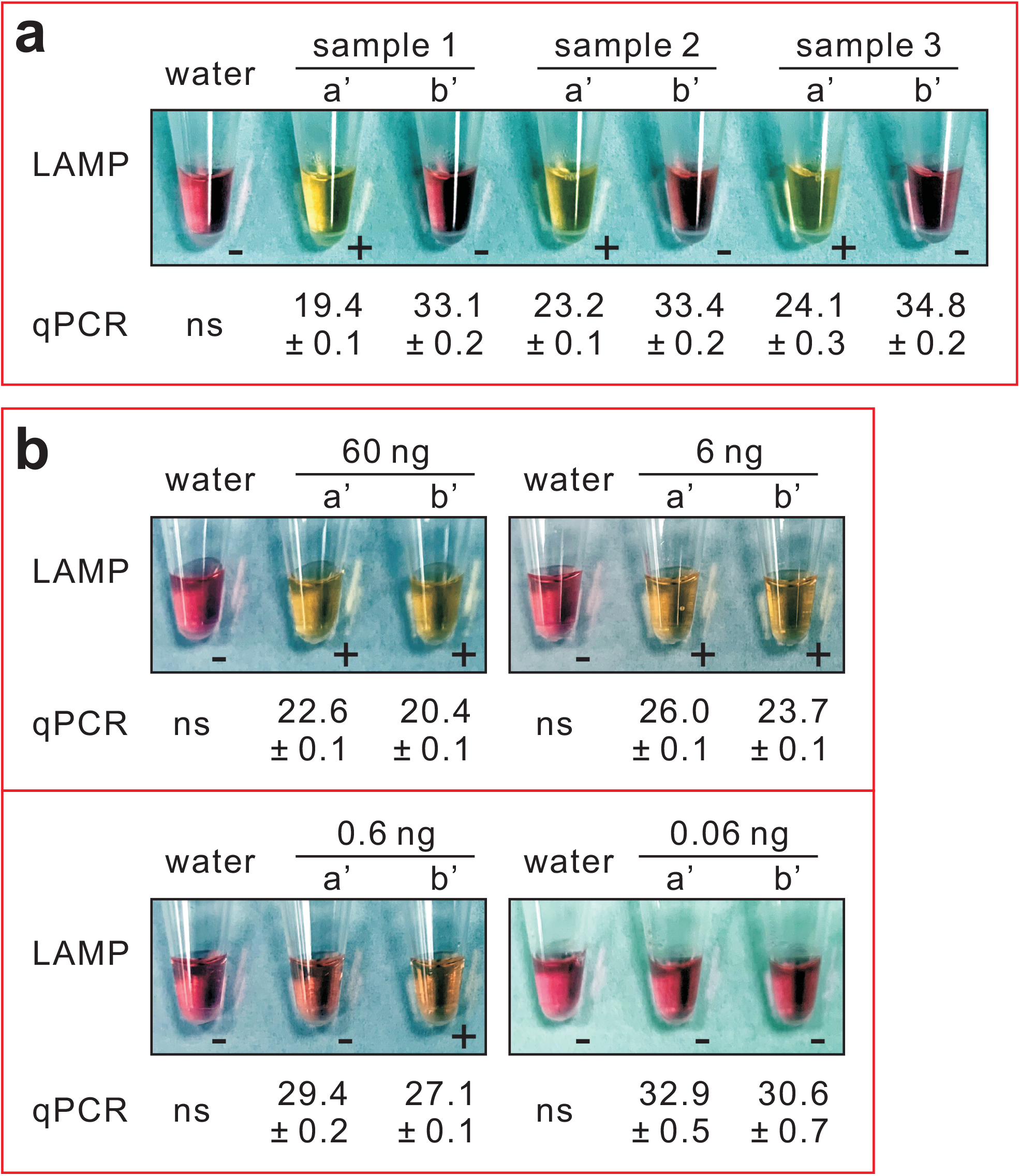
LAMP and qPCR test on infected wheat samples. + and -, positive and negative LAMP reaction, respectively. ns, no Cq value. The Cq values of qPCR are shown as mean ± standard deviation (n=3). (a) Test on DNA from three wheat seed samples. a’, from seed surface. b’, from washed seeds. (b) Test on DNA from infected leaves. a’ and b’, two leaf samples.

## Discussion

Currently there is no published PCR-based protocol that can differentiate Xtu and Xtt. In this study, F11/R11/P11 was designed based on an Xtt gene. This gene is also present in the Xtu genome with three nucleotides differing to its Xtt homolog. Thus, F11/R11/P11 can be used for detection of both Xtu and Xtt but not for differentiation of the two pathovars. The desire to have a PCR-based technique to differentiate Xtu from Xtt has been intensified by the observation that Xtt was generally nonpathogenic to wheat and thus detection of Xtt on wheat seed surface was believed not to be a severe concern (Sapkota et al. 2018). However, Curland et al. (2018) reported that most Xtu and Xtt strains collected from the Upper Midwestern United States could cause symptoms on both wheat and barley, even though their virulence was generally correlated with their original host. The target of F11/R11/P11, GenBank accession number SCB06003, may potentially be used to differentiate Xtu from Xtt. At the locus of SCB06003, there are three single nucleotide polymorphism (SNP) between the sequences in Xtt and Xtu strains. These SNPs can be utilized to design RNase H-dependent PCR (rhPCR) primers (Yang et al. 2018). In addition, one of the three SNPs creates a *Bgl* I cutting site in Xtu, which is absent in Xtt. Thus, restriction fragment length polymorphism (RFLP) is another potential method to differentiate Xtu and Xtt by utilizing this gene. However, both RFLP and rhPCR are based on conventional PCR and their sensitivities would be compromised when compared to qPCR.

It has not escaped our notice that some primer pairs in our 27 primer pair list (Table S1) may be good candidates for designing qPCR primers for Xtt detection. For example, the sequence information suggests that the target gene and primer sequences of primer pair F13/R13, are present only in Xtt. With the availability of confirmed Xtt strains, we will test these primer pairs in future studies.

It was not our intention to make unfavorable comparison between LAMP and qPCR. Rather, we desired to use the established LAMP method as a baseline from which to evaluate the efficiency of our qPCR system. In general LAMP should be more sensitive than conventional PCR and as sensitive as qPCR, which has been demonstrated in the diagnosis of many other pathogens (Tiberini et al. 2019; Wang et al. 2019; Sarkes et al. 2019). The LAMP targets an intergenic region (between GenBank accession number SCB03893 and SCB03894) and is specific to pv. *undulosa*, pv. *translucens* and pv. *secalis*. This LAMP system is very useful and has been used in our lab for diagnoses of wheat seed samples. The relative low sensitivity of LAMP in our lab, might be due to the fact that Langlois et al. (2017) and the current study used different LAMP chemicals.

In the sensitivity test, when DNA dilutions were used as templates, probe-based qPCR had efficiency of 0.99, which is in the normal range of 90-105% (Johnson et al. 2013). When DNA from cell dilutions were used, the efficiency was 1.28. We attributed the usually high efficiency to the PCR inhibitors present in the bacterial cells. Greater numbers of bacterial cells give higher concentrations of inhibitors in the resultant DNA.

From this study we report that the probe-based qPCR was more specific than SYBR Green-based qPCR. The probe P11 did not create new specificity, but rather, it revealed the existing specificity. The Cq values of the SYBR Green-based qPCR on non-target DNA indicated that the primer pair F11/R11 could produce non-specific amplification. On this non-specific amplification, SYBR Green generated signals but the probe did not. Thus, the probe-based qPCR was more specific than the SYBR-Green based qPCR, which is an observed result but not a norm. Indeed a plant diagnostic lab would prefer SYBR Green-based qPCR than probe-based qPCR, but only if no nonspecific amplification is produced by the primers. This is because the probe-based method is far more expensive than SYBR Green-based qPCR.

The low detection limit of our probe-based qPCR system is 60 fg DNA per reaction. According to Falahi et al. (2017), the genome of an Xtu strain is 4.6 Mb, with a calculated weight per DNA molecule of 4.8 fg (http://www.calctool.org/CALC/prof/bio/dna). Thus the qPCR could detect the target from DNA of 60/4.8 = 12.5 cells, given the assumption that 100% DNA of the cells was extracted. However, this was impossible in practice, as indicated by our result that 1,000 bacterial cells was the low limit of detection. The limit might be lower than 1000 cells, but likely more than 100 cells, as indicated by our test on DNA from 100-cell samples. These results emphasized that DNA extraction had significant influence on the sensitivity of a qPCR diagnosis system (Yang et al. 2021). Thus we highly recommend that DNA extraction efficiency should be considered when developing and advocating new PCR-based methods. When claiming the sensitivity, whenever it is possible, one should indicate the lowest number of spores or cells from which the DNA in one positive reaction was derived. This should also be considered when interpreting results from previously published PCR-based detection or quantification studies.

Considering sample preparation, DNA extraction and qPCR as an integrated diagnostic system, we could calculate how many bacterial cells the system can detect. The low limit of detection from cell suspension was 1,000 cells, which meant 40 cells per reaction. In the seed testing protocol we developed, one DNA sample contained DNA from the surface of 25 seeds, thus one seed in one qPCR reaction. Therefore, our qPCR system can detect the bacteria if one seed carries ≥ 40 bacterial cells, which is comparable with the results of Schaad and Forster (1985) who reported that seed lots with fewer than 1,000 cfu/mL in seed washing (approximately 42 cfu/seed) were unlikely to develop disease under irrigated growing conditions. It is worth mentioning that 42 cfu might be far more than 42 bacterial cells.

Taken together, we concluded that our qPCR system is useful for BLS seed testing and identification of the pathogen from infected plants. The system is specific to the pathovars *translucens* and *undulosa* but can’t differentiate the two pathovars. The sensitivity is on or below the minimal level of the pathogen that could cause an epidemic.

## Notes

### Competing Interest Statement

The authors have declared no competing interest.

