## Supplementary material for "Detection of *Xanthomonas translucens* pv. *undulosa* from wheat by quantitative PCR": Table S1

**Table S1.** All primers designed for *Xanthomonas translucens* detection and the presence/absence of their homologs

| | Primer | Target gene | Expected size (bp) | pv. *translucens* | pv. *undulosa* | pv. *cerealis* | Other |
|---|---|---|---|---|---|---|---|
| F1 | CCCGACTATATGGCGGCAGT | SCB03388.1 | 110 | Y* | N | N | N |
| R1 | AACGCGTGGATCTGGTGGAA | | | Y | Y | N | Y |
| F2 | CGCCGATGCACGACACG | SCB03507.1 | 108 | Y | Y | N | Y |
| R2 | CTCTGCCACCAGGACCCTC | | | Y | Y | N | Y |
| F3 | CTGTACGGTCTGCGGTGACA | SCB03750.1 | 84 | Y | N | N | N |
| R3 | CAGTTCAAGGTATTCGCCGGG | | | Y* | N | N | N |
| F4 | TCAATCAACAGCGCCAGCAG | SCB03852.1 | 100 | Y | Y | N | N |
| R4 | CGCAAGGCTTGACCCATGAC | | | Y | N | N | N |
| F5 | AGGGGCTTTTGTTAGTTGCTCT | SCB03898.1 | 85 | Y | N | N | N |
| R5 | GGCGGAAAGCGTCTCCGATA | | | Y | N | N | N |
| F6 | GCCGAGAACGCCAATTCCAA | SCB04443.1 | 80 | Y | Y | Y* | N |
| R6 | CTCGGACAGCCGGTCATGG | | | Y | Y | N | Y |
| F7 | GCAGTTCCATGTGGTCACGC | SCB04822.1 | 80 | Y | Y | N | N |
| R7 | CTACCTGGTCAGCCCCGATG | | | Y | Y | N | Y* |
| F8 | CATTGTTGTCGCAGGTCCCG | SCB05767.1 | 109 | Y | N | N | N |
| R8 | CCGGCTTTCTCTTCCACCGT | | | Y | N | N | Y* |
| F9 | AGGGATGGCTGAAGTCTGGC | SCB05860.1 | 87 | Y* | N | N | N |
| R9 | ACGACAAAAGTCGTATTCCGCTG | | | Y* | N | N | N |
| F10 | TTAGGCGTATGGCGGGGTG | SCB05899.1 | 110 | Y | Y | N | N |
| R10 | TGTACTGGGATCGCGCCAC | | | Y | Y | N | N |
| F11 | AATTGCACTGGCTGTCGCTG | SCB06003.1 | 80 | Y | Y | N | N |
| R11 | GGCAAGCGCCAAGAAGGAAA | | | Y | Y | N | N |
| F12 | CCATAGCTGCCAACCTGCCG | SCB06254.1 | 89 | Y | Y | Y* | N |
| R12 | GCGCGCCCTGCGTCAAG | | | Y | N | N | Y |
| F13 | CTCGCAGAAATCCAGGGCCA | SCB06379.1 | 81 | Y | N | N | N |
| R13 | GACTACACGCTGGCTTTGCC | | | Y | N | N | N |
| F14 | ACAAAGCCACATGGAGCGTA | SCB06394.1 | 98 | Y* | N | N | N |
| R14 | CCTGCGACTGGGCTGAAGTA | | | Y | N | N | N |

**Table S1.** All primers designed for *Xanthomonas translucens* detection and the presence/absence of their homologs (Cont.)

| | Primer | Target gene | Expected size (bp) | pv. *translucens* | pv. *undulosa* | pv. *cerealis* | Other |
|---|---|---|---|---|---|---|---|
| F15 | CGGACGCCCGATATGTTCCT | SCB06409.1 | 104 | Y* | N | N | N |
| R15 | TTGCTGACGATGCCAGTGGA | | | Y* | N | N | N |
| F16 | TATCGACACGCTGCGGGAAG | SCB06794.1 | 107 | Y | Y | Y* | N |
| R16 | CCGGATCCACCTCCAGGAAC | | | Y | N | N | N |
| F17 | CAGACAGCTCTCGGGCACTT | SCB03888.1 | 87 | Y | N | N | N |
| R17 | GCGACCGAATTGGCAAACCA | | | Y* | N | N | N |
| F18 | ATAGGTCCACTGGTCGCTGC | SCB04026.1 | 99 | Y | Y | Y* | N |
| R18 | ATCGAGCCATTCCAGGCGAA | | | Y | N | N | N |
| F19 | CCGATCCGCTCGACGACTAC | SCB04234.1 | 81 | Y | Y | Y* | Y |
| R19 | GCCAACCGGACGTGGTCAA | | | Y | N | N | N |
| F20 | TCAGGTCATCGCCCAGGATT | SCB02878.1 | 88 | Y | Y | N | N |
| R20 | TGGCCCGGAGAACTTTTCCTT | | | Y | N | N | N |
| F21 | GTCGAGCAGCTTGAACACGG | SCB03813.1 | 83 | Y | Y | N | Y* |
| R21 | CCATCGTGTTCGGGGTCAAC | | | Y | Y | N | Y |
| F22 | GCGAGATCCCTTGTACGGCT | SCB04824.1 | 86 | Y | Y | N | N |
| R22 | TTCTGTTTCGTTGGCAGCCC | | | Y | N | N | Y |
| F23 | CGGCACCTGGATGACGATCT | SCB05509.1 | 86 | Y | N | N | Y |
| R23 | TAAGCGGCGTAGCGATCGAA | | | Y | N | N | N |
| F24 | GCATATCCACGCTCGCCAAC | SCB06089.1 | 91 | Y | N | N | N |
| R24 | TACTTCCGGCGCTCAAGGAG | | | Y | N | Y* | N |
| F25 | TCGGGATGCAGCAGATCGTC | SCB06297.1 | 80 | Y | Y | Y* | Y |
| R25 | GCCGCATTGCAGGCATGG | | | Y | Y | N | Y |
| F26 | AAGACGACGACCAGCACGAT | SCB06299.1 | 107 | Y | Y | N | Y* |
| R26 | GAGCGCTGGCTGGAAGAACA | | | Y | N | N | Y |
| F27 | CCATGGGGTTGCAGGCGAG | SCB06730.1 | 98 | Y | N | N | Y* |
| R27 | CGTTGTCCGGCACCACGAT | | | Y | Y | N | Y |

*From one strain only
